# Stochastic spin selection in the mechanism of semiquinone formation at the ubiquinol oxidation Q_o_ site of cytochrome *bc*_1_

**DOI:** 10.64898/2026.05.22.727096

**Authors:** Łukasz Bujnowicz, Rafał Pietras, Anna Wójcik-Augustyn, Artur Osyczka, Marcin Sarewicz

## Abstract

Cytochrome *bc*_1_ is one of the key enzymes of biological energy-conserving systems. In its catalytic Q cycle, the central reaction is the oxidation of quinol (QH_2_), upon which electrons are directed to two separate cofactor chains. The molecular mechanism of this reaction remains elusive. The canonical model, assuming a sequence of reactions dictated by the equilibrium redox midpoint potentials of cofactors (the 2Fe2S cluster and heme *b*_L_), has recently been challenged by a new model of EB derived from quantum mechanical (QM) calculations – EMET (EMergent Electron Transfer) (https://doi.org/10.1021/acsomega.5c13233). These two models predict fundamentally different microstates of the enzyme in which semiquinone (SQ) is formed in the catalytic site (Q _o_) and also predict different lowest-energy configurations. Here, we test these predictions using EPR spectroscopy on highly concentrated preparations of isolated bacterial cytochrome *bc*_1_. We detect SQ spin-coupled to the reduced 2Fe2S cluster (2Fe2S^red^), whose population markedly exceeds that of reduced heme *b*_L_ and forms exclusively in sites containing oxidized heme. We also identify that the lowest-energy configuration corresponds to the state with reduced heme *b*_H_ (adjacent to heme *b*_L_), oxidized heme *b*_L_ and SQ-2Fe2S^red^. These two features are precluded by the canonical model but are consistent with EMET. We conclude that EMET, unlike the canonical EB model, satisfactorily describes the occurrence of stochastic, spin-selective processes that result in electron stoichiometry among hemes *b*, the 2Fe2S cluster, and SQ at Q_o_ that are observed spectroscopically.

## Introduction

Cytochrome (cyt.) *bc*_1_, a member of the cytochrome *bc* (Rieske/b) family [1], catalyzes oxidation of quinols (QH_2_) and reduction of cyt. *c*, contributing to the proton-motive force driving ATP synthesis [2–4]. Cytochromes *bc* share a conserved homodimeric architecture and nearly identical arrangements of redox cofactors and quinol/quinone binding sites (Q_o_ and Q_i_), supporting a broadly conserved catalytic mechanism across species, despite differences in subunit composition or auxiliary cofactors in complexes such as cyt. *b*_6_*f* (Fig. 1) [5–15].

**Figure 1.**
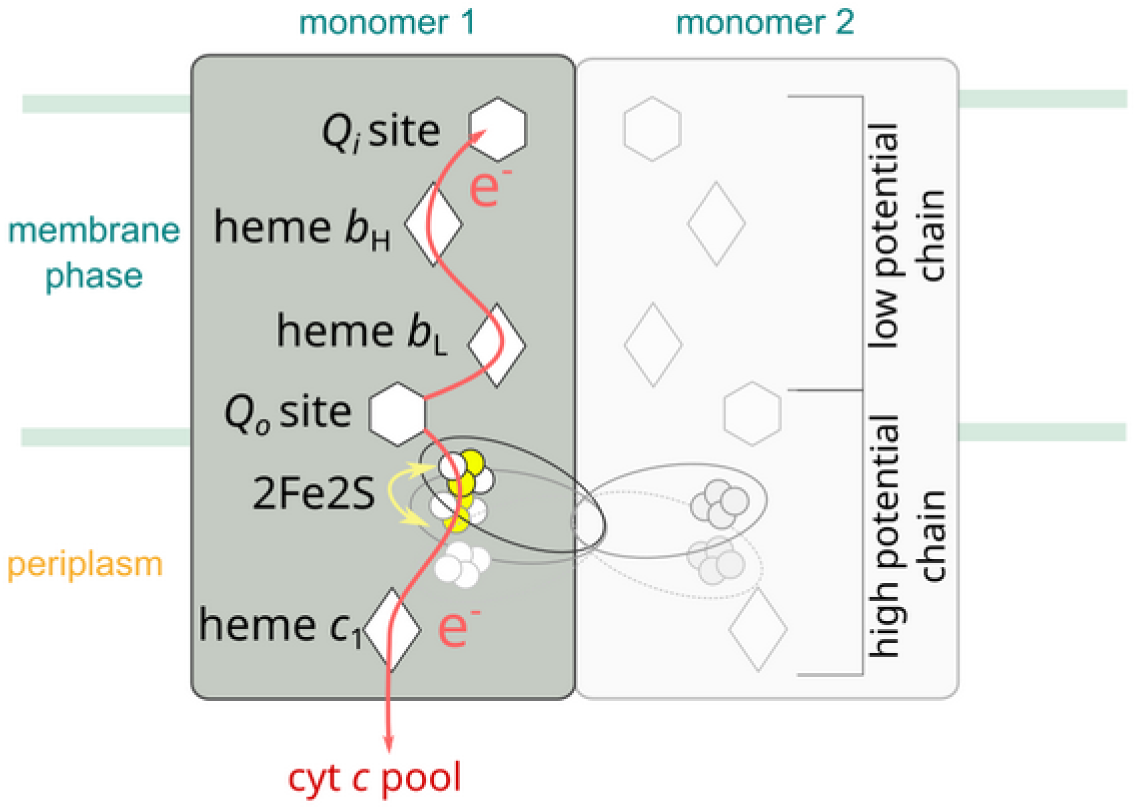
Schematic representation of the cytochrome *bc*_1_ dimer. The enzyme is embedded in the lipid membrane (green lines) and consists of two identical monomers (1 and 2). For clarity, only monomer 1 is shown in full color. Each monomer contains two quinone-binding sites (hexagons): the Q_o_ site, where electron bifurcation occurs, and the Q_i_ site, where quinone undergoes two-step reduction. Electrons released from QH_2_ at the Qo site are split into two branches. The low-potential chain comprises heme *b*_L_ and *b*_H_ (white rhombi) and directs electrons toward the Q_i_ site. The high-potential chain includes the Rieske 2Fe2S cluster (yellow circles), located within the mobile ISP-HD (ellipse), which transfers electrons from the Q_o_ site to heme *c*₁ and subsequently to the cytochrome *c* pool. The red arrows depict electron-transfer steps. The yellow double-headed arrow indicates the movement of the ISP-HD around its pivot point.

The catalytic mechanism of QH_2_ oxidation, known as the Q cycle, was originally proposed by Mitchell and subsequently modified by Crofts et al. [16,17]. A central feature of this mechanism is electron bifurcation (EB), in which two electrons derived from QH_2_ are transferred to two spatially separated acceptors: heme *b*_L_ and the Rieske 2Fe2S cluster (2Fe2S). In this way the route for two electrons, originating from QH_2_, diverges into two separate chains: the low- and the high-potential chain (Fig. 1). In the low-potential chain the electrons are transferred from heme *b*_L_ to heme *b*_H_ and finally to the Q_i_ site, where quinone (Q) is reduced, while in the high-potential chain the electrons are transported *via* the mobile head domain of the iron-sulfur protein (ISP-HD) from the Q_o_ site to heme *c*_1_, which mediates electron transfer (ET) to water-soluble cyt. *c* [18–24]. The efficiency of EB depends critically on the coupling of electron and proton transfer (PT) during QH_2_ oxidation.

Until recently, quinone-based EB has been described using equilibrium redox potentials (*E*_m_) of 2Fe2S and heme *b*_L_. We refer to this as canonical QBEB [25–30]. It proposes that QH_2_ is first oxidized by the high-potential 2Fe2S cluster to produce a semiquinone (SQ) and then SQ is oxidized by the low-potential heme *b*_L_. These two steps are considered uphill and downhill reactions, respectively. Because the SQ/QH_2_ and Q/SQ couples are thought to have a relatively high and low redox potential, respectively, SQ is considered unstable and would not accumulate as long as heme *b*_L_ remains oxidized [28,30]. This is one of the most important conclusions drawn from canonical QBEB. This conclusion leads to expectation that, if SQ ever occupies the Q_o_ site, it does so under conditions that keep heme *b*_L_ in a reduced state [31]. This imposes the risk of short circuit reactions initiated by ET from heme *b*_L_ back to the Q_o_ site, resulting in fast cyt. *c* reduction, despite lack of proton translocation across the membrane [3]. As such reactions need to be suppressed to secure energetic efficiency of cyt. *bc*_1_, various suppression mechanisms have been proposed, but none has been proven experimentally [28,32–40]

The conditions favoring a maintenance of heme *b*_L_ in reduced state were thus considered as a premise for detection of SQ in the Q_o_ site. As blocking the Q_i_ site by antimycin has long been known to favor heme *b*_H_ in the reduced state, it was assumed that that this inhibitor indirectly favors a state in which heme *b*_L_ is also reduced, thereby increasing a probability for detection of SQ [32,41–43]. In fact, weak EPR radical signals at g = 2 have been reported, although these studies did not establish the redox state of heme *b*_L_ in the presence of SQ [30,44–47]. Furthermore, one of those reports showed paramagnetic interaction between heme *b*_L_ and SQ, which requires heme *b*_L_ to be oxidized, thereby contradicting the expectations of canonical QBEB [46].

In our studies on *Rhodobacter capsulatus* cyt. *bc*_1_, we also detected the SQ at the Q_o_ site in the antimycin-inhibited enzyme. However, in this case the amount of the signal was substantial, which was unexpected considering the high instability of SQ postulated by the canonical QBEB mechanisms [48–51]. Interestingly, most of the SQ fraction was detected as a rhombic EPR signal (SQ–2Fe2S^red^) resulting from the spin-coupling between SQ and the reduced 2Fe2S cluster, instead of the expected radical signal [51]. In a mutant with the changed axial ligand of heme *b*_L_, the SQ-2Fe2S^red^ signal was not detectable suggesting involvement of the heme in the process of SQ-2Fe2S^red^ formation [52]. On the other hand, substantial amounts of SQ-2Fe2S^red^ signal was detected in non-inhibited cyt. *b*_6_*f* from spinach [50]. These observations seemed difficult to explain within the framework of canonical QBEB, which predicts either rapid oxidation of SQ by oxidized heme *b*_L_ or, alternatively, its reduction to QH_2_ by ET from reduced heme *b*_L_ in the absence of gating.

To understand the origin of the EPR-detectable species, we recently performed quantum mechanical (QM) calculations on large cluster models of cyt. *bc*_1_ that simultaneously considered both electron acceptors from QH_2_. These calculations revealed an alternative mechanism of electron bifurcation, which we referred to as EMET (EMergent Electron Transfer). According to EMET, the QH₂ oxidation requires presence of oxidized 2Fe2S and the reaction begins with ET from QH_2_ to heme *b*_L_. The second electron from SQ to oxidized 2Fe2S occurs only after prior ET from heme *b*_L_ to heme *b*_H_ and/or proton release from the Q_o_ site [53]. This sequence of ET results from cooperation between two redox cofactors directly involved in the oxidation of QH₂ at the Q_o_ site and can be modeled only when both cofactors are treated at the QM level. However, if heme *b*_L_ cannot pass the electron to heme *b*_H_ (as in the presence of antimycin), SQ can be stabilized, which would manifest as ferromagnetic spin coupling with reduced 2Fe2S (2Fe2S^red^), detectable by EPR. On the other hand, the antiferromagnetic coupling between SQ and 2Fe2S^red^ is unstable and tends to relax back to QH_2_ and oxidized 2Fe2S. This mechanism emerged when cluster models used for QM calculations included both immediate electron acceptors simultaneously, rather than splitting the bifurcation into two independent steps as done in previous studies [54–57]. The unequal stability of ferro- and antiferromagnetic SQ–2Fe2S^red^ coupling predicts that full Q_o_-site occupancy by SQ–2Fe2S^red^ is statistically impossible, yielding an expected 1:1 partition between productive SQ–2Fe2S^red^ formation and relaxation back to QH_2_ due to stochastic spin selection. One of the resulting configurations recognized by theoretical studies is characterized by reduced heme *b*_H_, oxidized heme *b*_L_, and SQ spin-coupled to 2Fe2S^red^.

Experimental detection of this particular configuration would be one of the crucial indicators in support of EMET. As mentioned above, previous reports showed that SQ–2Fe2S^red^ is detected along with oxidized heme *b*_L_. Although this constitutes a strong argument supporting the conclusions drawn from theoretical studies, it cannot be quantitatively demonstrated that the SQ–2Fe2S^red^ species is present in the exact same monomer of cyt. *bc*_1_ in which heme *b*_L_ is oxidized. This is due to the limited sensitivity of previous CW EPR measurements. Typically, the amount of SQ–2Fe2S^red^ generated in those experiments was much lower than 50 % of total Q_o_ sites. Additionally, the g_z_ EPR transition of oxidized heme *b*_L_ is very weak, and the uncertainty in determining the reduced heme *b*_L_ fraction reaches ∼20–30 % of the total heme. Because the fraction of SQ–2Fe2S^red^ is comparable to this uncertainty, it is not possible to rule out the possibility that SQ–2Fe2S^red^ is generated in the same monomers containing reduced heme *b*_L_.

In this study, we analyzed isolated *R. capsulatus* cyt. *bc*_1_, both wild type (WT) and the mutant with severely restricted ISP-HD movement (+2Ala) [19], under defined redox conditions to characterize the electronic configuration of individual Q_o_-site monomers. Our goals were to identify conditions that promote formation of the SQ–2Fe2S^red^ state, quantify the redox balance among cofactors involved in EB, and compare these results with predictions from the EMET mechanism. We prepared highly concentrated samples frozen at multiple equilibrium and non-equilibrium redox states to directly and quantitatively detect all relevant cofactors. Additionally, pulsed EPR distance-dependent relaxation measurements allowed us to determine the distance between oxidized heme *b*_L_ and SQ–2Fe2S^red^, demonstrating that SQ occupies a position consistent with the stigmatellin-binding position [57], and that this state forms exclusively in monomers containing oxidized heme *b*_L_. Finally, we show that the probability of SQ–2Fe2S^red^ formation in monomers with antimycin-occupied Q_i_ sites equals 0.5, consistent with the stochastic spin selection predicted by the EMET mechanism.

## Materials and methods

Equine cytochrome *c*, decylubiquinone (DB), antimycin A, sodium borohydride, sodium dithionite, potassium ferricyanide (PFC) and other reagents were purchased from Sigma-Aldrich. *n*-Dodecyl-β-D-maltoside (DDM) detergent was purchased from Anatrace. DB was suspended in DMSO and reduced to its hydroquinone form (DBH₂) using hydrogen gas released from an acidic aqueous solution of sodium borohydride in the presence of platinum on carbon.

Cyt. *bc*₁ from wild-type *Rhodobacter capsulatus* grown under semiaerobic conditions was isolated according to the procedure described in ref. [59] but with no glycerol added using Bio-Rad NGC FPLC system. The isolated proteins were dissolved in 50 mM HEPES buffer, pH 8.0, containing 100 mM NaCl, 1 mM EDTA, and 0.01 % DDM and subsequently antimycin A, cyt. *c*, and PFC were added to the solution. A 50 µL aliquot of this mixture was rapidly mixed with 150 µL of DBH₂ solution in the same buffer. After a short incubation (3 s or 8 s), the samples were frozen in an EPR measurement tube. The final concentrations were as follows: antimycin A, 345 µM; cyt. *c*, 96 µM; PFC, 575 µM; DBH₂, 750 µM. The concentration of +2Ala mutant and WT cyt. *bc*_1_ was 109 and 96 µM, respectively.

Continuous wave (CW) and pulsed EPR measurements were performed on the same samples at X band. After measurement of a sample prepared at given conditions the sample was thawed and allowed to reach the redox equilibrium. Then, the sample was frozen again and the EPR measurements were repeated. To fully oxidize or reduce the samples 2 µL of 300 mM PFC solution or few grains of sodium dithionite was added, respectively.

The EPR measurements were performed using a Bruker Elexsys E580 spectrometer equipped with a ColdEdge, the stinger closed-cycle helium cryocooler. CW EPR spectra were measured with the use of a Bruker SHQ4122 resonator cavity fitted with an Oxford Instruments ESR900 cryostat. The measurement parameters for the iron–sulfur cluster spectra were as follows: temperature 20 K, center field 3607 G, sweep width 927 G, modulation amplitude 16 G, modulation frequency 100 kHz, time constant 82 ms, conversion time 41 ms, sweep time 41 s, and microwave power 2 mW. Parameters for measurement of hemes *b* were: temperature 10 K, center field 1918 G, sweep width 1062 G, modulation amplitude 16 G, modulation frequency 100 kHz, time constant 20 s, conversion time 10 ms, sweep time 41 s, and microwave power 2 mW.

The electron spin echo (ESE) decay curves were registered at magnetic field positions corresponding to g = 1.94 and g = 1.85 using Hahn echo sequence: π/2 – (100 ns + *t)* – π – (100 ns + *t)* – echo. The pulse lengths were 20 ns and 40 ns, respectively and the interpulse delay *t* was varied from 0 to 3000 ns. The repetition time of the pulse sequence was 500 μs, with 40 averages per point, a signal gain of 42 dB, and 20–60 scans per decay curves. Measurements were performed at the following temperatures: 10 K, 12 K, 14 K, 16 K, 17.5 K, 19.5 K, and 21 K.

## QM models and methods

Large cluster models used in the QM calculations were constructed based on the selected optimized intermediate states described in our previous work [53], which were extended to include the optimized heme *b*_H_ fragment in its reduced state. They consist of 546 atoms and include: heme *b*_H_, heme *b*_L_, ubiquinone/semiquinone (with tail truncated to one isoprene unit), 2Fe2S, two water molecules, eight residues from ISP (C133, H135, C138, C153, C155, H156, S158, Y160) and sixteen residues from cyt. *b*: twelve applied previously in model of Q_o_ site [53] (R94, H97, Y147, M154, H198, H276, D278, N279, V293, P294, E295, Y302) and four from direct surrounding of heme *b*_H_ (W45, H111, H112, R114). The following residues: C133, H135, C138, C153, S158, Y160 from ISP and W45, R94, H97, Y147, M154, H198, H111, H212, R114, H276, Y302 from cyt. *b* were included with their -NH and -CO moieties of the protein backbone substituted by hydrogen atoms. The protein fragments C155–H156 (ISP) and D278–N279, V293–P294–E295 (cyt. *b*) were included along with the peptide bonds connecting them, with the corresponding carbonyl and amino groups of the protein backbone replaced by hydrogen atoms. Protonation states of the applied residues were validated by Propka 3.1 and pypKa 2.10.0 [60–62] programs. The QM models were assembled from the optimized intermediate described previously [53] for the antimycin-inhibited enzyme (P3, P2 or P1^Y^) and optimized small model consisting of reduced heme *b*_H_ with its ligands (H111 and H212) and interacting residues (W45, R114). Small model of heme *b*_H_ was optimized with the procedure applied previously: the DFT/B3LYP method enriched with Grimme’s D3 dispersion correction with Becke–Johnson damping and combined with the double-ζ basis set def2-SVP [63–65]. The large models assembled from fragments optimized separately were applied to estimate the energies of different electronic configurations using the DFT/B3LYP-D3/ def2-SVP. The QM calculations were performed using the Gaussian 16 program [66].

## Results and Discussion

### CW EPR spectra under non-equilibrium redox conditions

In Fig. 2 the (CW) EPR spectra measured for WT cyt. *bc*_1_ at different redox conditions are shown. The spectra were registered at two different ranges of the magnetic field. The first of the ranges covered the EPR transitions, originating from 2Fe2S^red^ and SQ spin coupled to 2Fe2S^red^, measured at 20 K. The second range involved the magnetic fields, where g_z_ transitions originating from the oxidized hemes *b* contribute to the EPR signals, measured at 10 K.

**Figure. 2.**
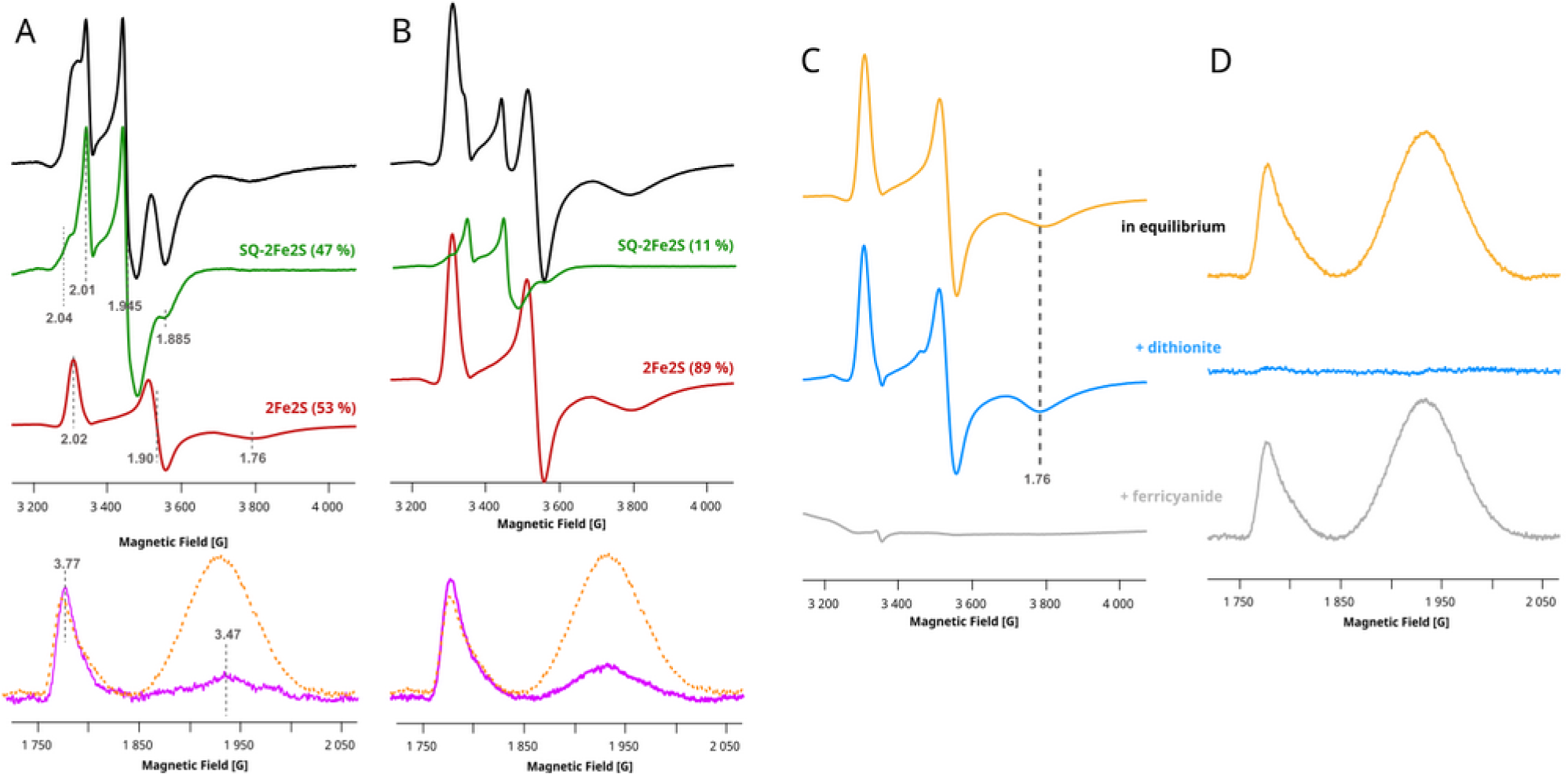
EPR spectra of the paramagnetic centers in WT cyt. *bc₁* measured under different redox conditions. **A)** Spectra measured for samples containing the enzyme frozen 3 s after mixing with substrates. The black spectrum represents the experimental spectrum being a sum of SQ–2Fe2S^red^ and 2Fe2S^red^ components, shown in green and red, respectively. Numbers in parentheses indicate the fractional contributions of each species to the black spectrum. The magenta line shows spectrum of the *g*_z_ transitions of hemes *b*_L_ and *b*_H_ (at *g* = 3.77 and 3.47, respectively), compared with the corresponding spectra recorded after reaching redox equilibrium (orange dotted lines). **B)** Same as in (A), but for samples frozen 8 s after mixing with the substrates. **C)** Spectra of 2Fe2S^red^ measured at redox equilibrium (orange), after reduction with dithionite (blue) and after oxidation with ferricyanide (gray). **D)** *g*_z_ transitions of hemes *b*_L_ and *b*_H_ recorded for the same samples as in (C). Vertical dashed lines indicate the *g*-values of the selected transitions.

For samples containing wild type (WT) cyt. *bc*_1_, mixed with the substrates (synthetic ubiquinol analog – decylubiquinol (DBH_2_) and the oxidized cyt. *c*) and frozen 3 s. after mixing, a spectrum reflecting of a sum of 2Fe2S^red^ and SQ-2Fe2S^red^ spectra is detected (Fig. 2A, black). After decomposition of the spectrum into these two components we obtain 47 and 53 % of the total cluster as SQ-2Fe2S^red^ and 2Fe2S^red^ (Fig. 2A green and red spectrum, respectively). We have been unable to generate SQ-2Fe2S^red^ exceeding 50 % of the total 2Fe2S cluster and this SQ-2Fe2S^red^ to 2Fe2S^red^ ratio has been found to be the highest. It is worth noting that the g_x_ transition of 2Fe2S^red^ is detected at g = 1.76, which is typical value observed for WT cyt. *bc*_1_ for which either Q_o_ site is empty or for which ISP-HD is not at the Q_o_ site, thus not interacting with the molecule bound at the site [67]. For the same sample, EPR spectra of the hemes *b* (Fig. 2A magenta) revealed that approximately 89 % of heme *b*_H_ (transition at g_z_ = 3.47) is in reduced state, which is expected for the enzyme with antimycin bound at the Q_i_ site. However, at the same time only 12 % fraction of hemes *b*_L_ is in the reduced state but this value is within the uncertanity of the qualitative EPR method (compare Fig. 2A orange dotted lines for fully oxidized hemes *b*).

For samples prepared in a similar way but frozen 8 s after mixing with the substrates, we observed a large decrease in the fraction of SQ–2Fe2S^red^ (to 11 % of the total cluster), along with a corresponding increase in 2Fe2S^red^ (Fig. 2B, black, green, and red, respectively). At the same time, heme *b*_H_ remained about 70 % reduced, while heme *b*_L_ remained oxidized (Fig. 2B magenta). Again, the g_x_ transition of 2Fe2S^red^ showed no interaction with the quinone molecule at the Q_o_ site as ISP-HD in isolated WT cyt. *bc*_1_ is predominantly in the remote position [23].

The sample frozen 8 s after mixing with the substrates was thawed and allowed to reach redox equilibrium within several minutes. The sample was then measured again, and we detected fully reduced 2Fe2S, with g_x_ showing no interactions with a quinone molecule at the Q_o_ site (Fig. 2C orange). In this case, hemes *b* were in fully oxidized state (Fig. 2D orange). The maximum reduction level of 2Fe2S^red^ and the maximum oxidation level of hemes *b* were obtained after reducing the same sample with dithionite or oxidizing it with ferricyanide, respectively (Fig. 2C and D, blue and gray, respectively). These control spectra allowed us to estimate the degree of reduction and oxidation for the samples frozen 3 and 8 s after mixing with the substrates. The results are summarized in Table 1.

**Table 1.** Fractions of the reduced cofactors and the fraction of SQ-2Fe2S^red^ for WT and 2Ala obtained after 3 or 8 sec after mixing with the excess of the substrates.

| Sample | 2Fe2S<sup>red</sup> | SQ-2Fe2S<sup>red</sup> | $b_L^{\text{red}^*}$ | $b_H^{\text{red}^*}$ |
| --- | --- | --- | --- | --- |
| WT 3 sec. | 53 | 47 | 12 | 89 |
| WT 8 sec. | 88.6 | 11.4 | 0 | 70 |
| 2Ala 8 sec. | 57 | 43 | 0 | 77 |

\* The fraction of reduced $b_L$ (or $b_H$ ) was estimated under the assumption of complete oxidation of the heme at equilibrium.

**Table 2.**
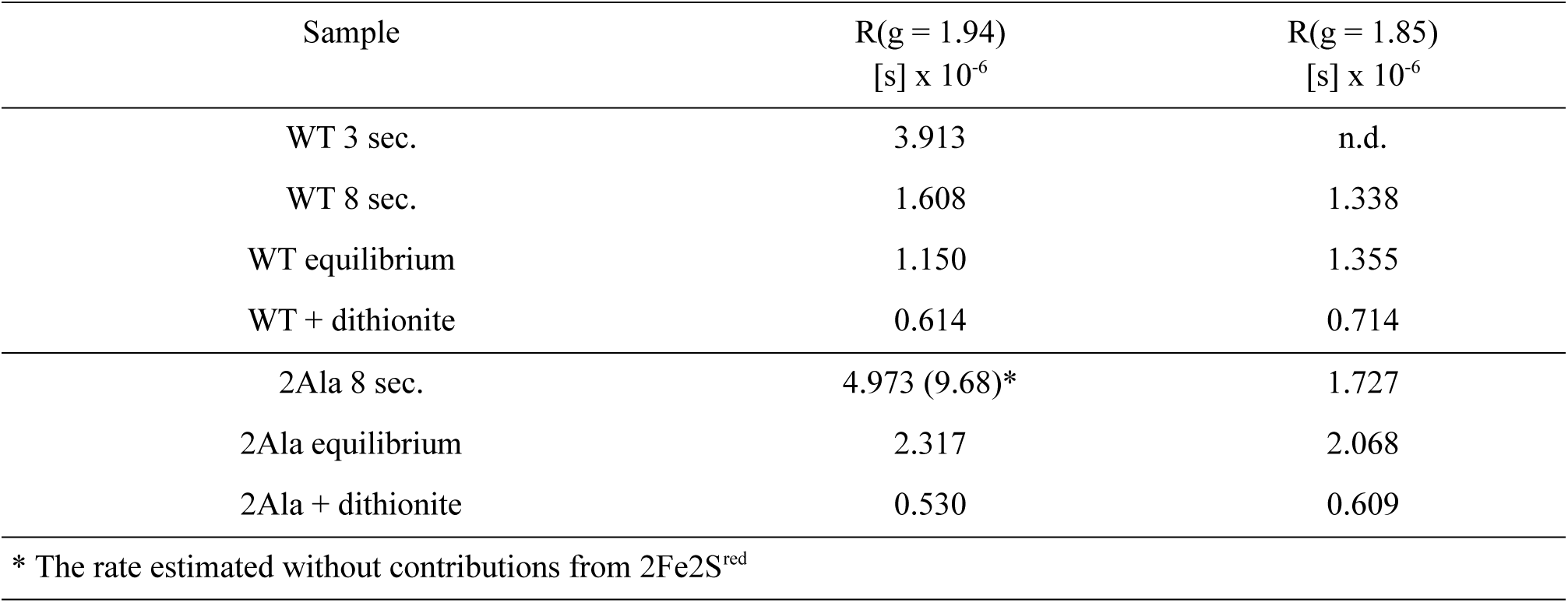
Phase relaxation rates (R) obtained for WT and 2Ala at 19.5K at g = 1.94 and g = 1.85.

| Sample | R($g = 1.94$)<br>[s] $\times 10^{-6}$ | R($g = 1.85$)<br>[s] $\times 10^{-6}$ |
| --- | --- | --- |
| WT 3 sec. | 3.913 | n.d. |
| WT 8 sec. | 1.608 | 1.338 |
| WT equilibrium | 1.150 | 1.355 |
| WT + dithionite | 0.614 | 0.714 |
| 2Ala 8 sec. | 4.973 (9.68)* | 1.727 |
| 2Ala equilibrium | 2.317 | 2.068 |
| 2Ala + dithionite | 0.530 | 0.609 |

\* The rate estimated without contributions from $2\text{Fe2S}^{\text{red}}$

Since formation of SQ-2Fe2S^red^ requires close contact between SQ and the cluster, we performed the same experiments as for WT cyt. *bc*_1_, but using the mutant which has the insertion of two Ala residues into the neck region of the iron–sulfur protein (+2Ala mutant) to see if SQ-2Fe2S^red^ fraction can be significantly increased. This mutation renders the ISP-HD to predominantly occupy the Q_o_ position when compared to WT [19,23,68]. In this case, the enzyme exhibited a much slower turnover rate due to the effect of the mutation [19,68–70]. The EPR spectra obtained for the mutant are shown in Fig. 3.

**Figure. 3.**
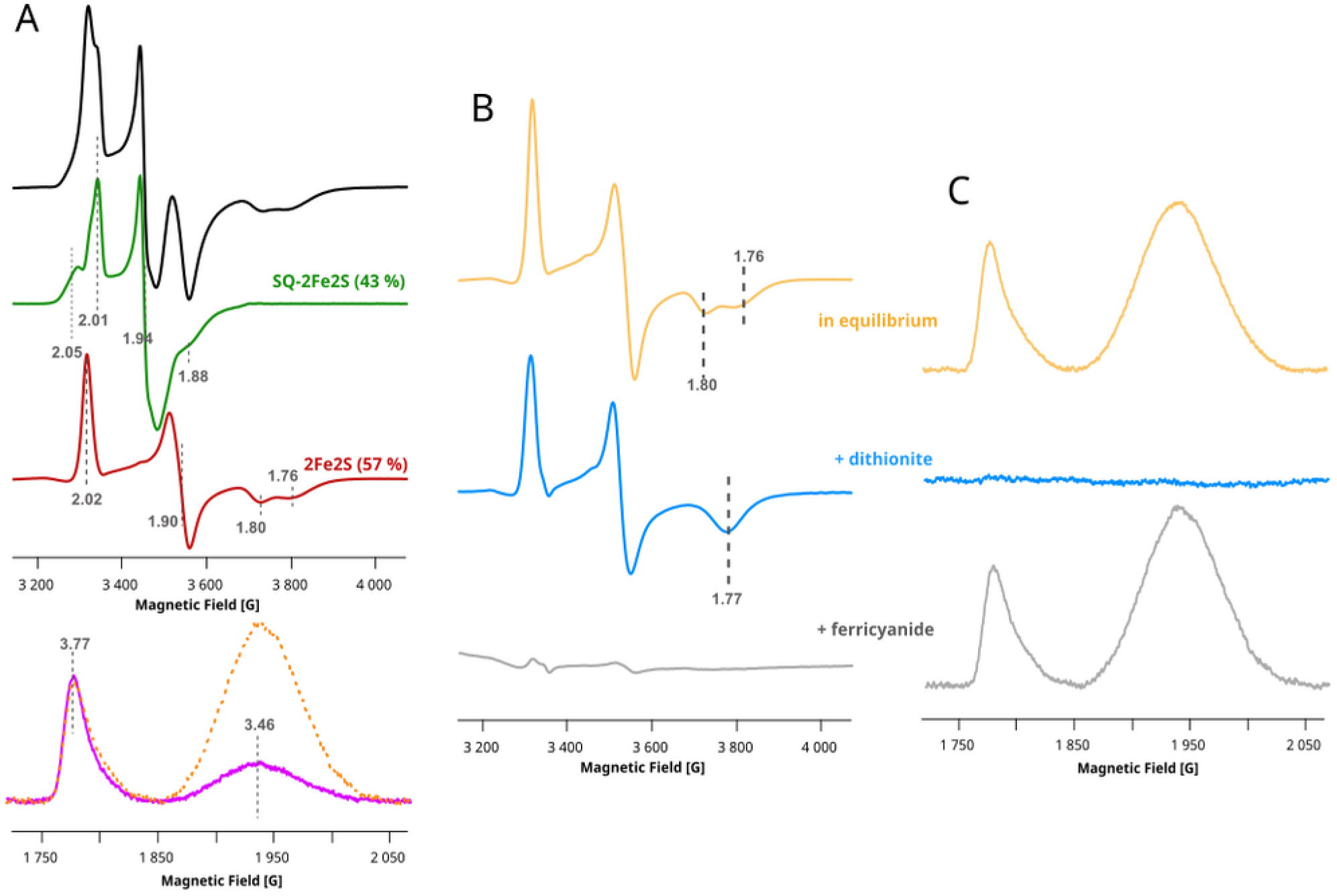
EPR spectra of the paramagnetic centers in +2Ala muntant of cyt. *bc₁* measured under different redox conditions. **A)** Spectra measured for samples containing the mutant enzyme frozen 8 s after mixing with substrates. The black spectrum represents the experimental spectrum being a sum of SQ–2Fe2S^red^ and 2Fe2S^red^ components, shown in green and red, respectively. Numbers in parentheses indicate the fractional contributions of each species to the black spectrum. The magenta spectrum shows the *g*_z_ transitions of hemes *b*_L_ and *b*_H_ (at *g* = 3.77 and 3.47), compared with the corresponding spectra recorded after reaching redox equilibrium (orange dotted lines). B) Spectra of 2Fe2S centers in the mutant cyt. *bc₁* measured at redox equilibrium (orange), after reduction with dithionite (blue), and after oxidation with ferricyanide (gray). C) *g*_z_ transitions of hemes *b*_L_ and *b*_H_ recorded for the same samples as in (C). Vertical dashed lines indicate the *g*-values of the selected transitions.

The samples of +2Ala mutant frozen 8 s after mixing with the substrates exhibit a mixture of 2Fe2S^red^ and SQ–2Fe2S^red^ components, contributing 57 % and 43 %, respectively (Fig. 3A, black, green, and red), similarly to WT frozen after 3 s. However, in contrast to WT cyt. *bc*_1_, we observed two distinct g_x_ transitions for 2Fe2S^red^, at g = 1.80 and g = 1.76. The presence of these two components in the cluster spectrum reflects interaction of the cluster with quinone occupying the Q_o_ site (g = 1.80). The transition at g = 1.76 suggests the absence of Q at the site or weaker interaction between the cluster and Q due to possible heterogenity of microconformations of the ISP-HD [71].

At the same time, the spectra showing the redox states of hemes *b* were very similar to those for WT cyt. *bc*_1_ frozen 8 s after mixing with the substrates. Heme *b*_L_ was fully oxidized, while heme *b*_H_ remained 77 % reduced (Fig. 3A magenta and Table 1).

After thawing the sample and allowing it to reach redox equilibrium, 2Fe2S became fully reduced while hemes *b* were fully oxidized (Fig. 3B and C, orange). Under these conditions, the +2Ala mutant still exhibited two distinct populations of the 2Fe2S cluster at the Q_o_site: one interacting with quinone (g_x_ = 1.80) and one not interacting with quinone (g_x_ = 1.76). The slightly increased amplitude of the g_x_ = 1.80 transition compared to spectra recorded before equilibrium reflects the higher fraction of Q formed during the enzymatic turnover.

In contrast, complete reduction of the quinone pool with dithionite yielded a homogeneous 2Fe2S ^red^ spectrum with a single g_x_ = 1.77 transition (Fig. 3B blue). This value is characteristic of the cluster interacting with QH_2_. A single g_x_ transition detected under reducing conditions indicates uniform occupation of the Q_o_. The disappearance of the fraction of the unoccupied Q_o_ sites (g = 1.76 transition) upon full reduction of quinone pool suggests that the reduced 2Fe2S cluster interacts more strongly with QH_2_ than with Q.

### Pulsed EPR measurements of the phase relaxation rates

The CW EPR spectra obtained for WT and +2Ala cyt. *bc*_1_ report the steady-state reduction levels of individual cofactors under different redox conditions. However, they do not provide information on the relative distances between transient SQ–2Fe2S^red^ species and the oxidized heme *b*_L_. If SQ–2Fe2S^red^ is indeed formed transiently at the quinone-binding Q_o_ site, in molecules where heme *b*_L_ is oxidized, the dipolar relaxation enhancement caused by the heme on the SQ–2Fe2S^red^ phase relaxation should be much stronger than for 2Fe2S^red^ [23,68]. This follows from the closer proximity of SQ to heme *b*_L_ within the Q_o_ pocket. Conversely, large-scale motion of the ISP-HD away from the Q_o_ site should weaken the dipolar enhancement by increasing the heme–cluster distance.

To quantify these effects, we measured ESE decay curves using the Hahn echo sequence for WT and +2Ala cyt. *bc*_1_ at two *g* positions: *g* = 1.94, which corresponds to the maximum of SQ–2Fe2S^red^, and *g* = 1.85, where the SQ–2Fe2S^red^ contribution is minor but the 2Fe2S^red^ signal remains detectable. Relaxation rates under the redox conditions shown in Figs. 2 and 3 were obtained by fitting a single-exponential function to the ESE decay traces.

Figure 4 shows the relaxation rates obtained for WT cyt. *bc*_1_ measured between 10 and 21 K at *g* = 1.94 (top) and 1.85 (bottom). When the enzyme is fully reduced by dithionite, heme *b*_L_ is diamagnetic and the dipolar interaction is absent. Under these conditions, the relaxation rate increases only slightly and monotonically with temperature (Fig. 4, black).

**Figure 4.**
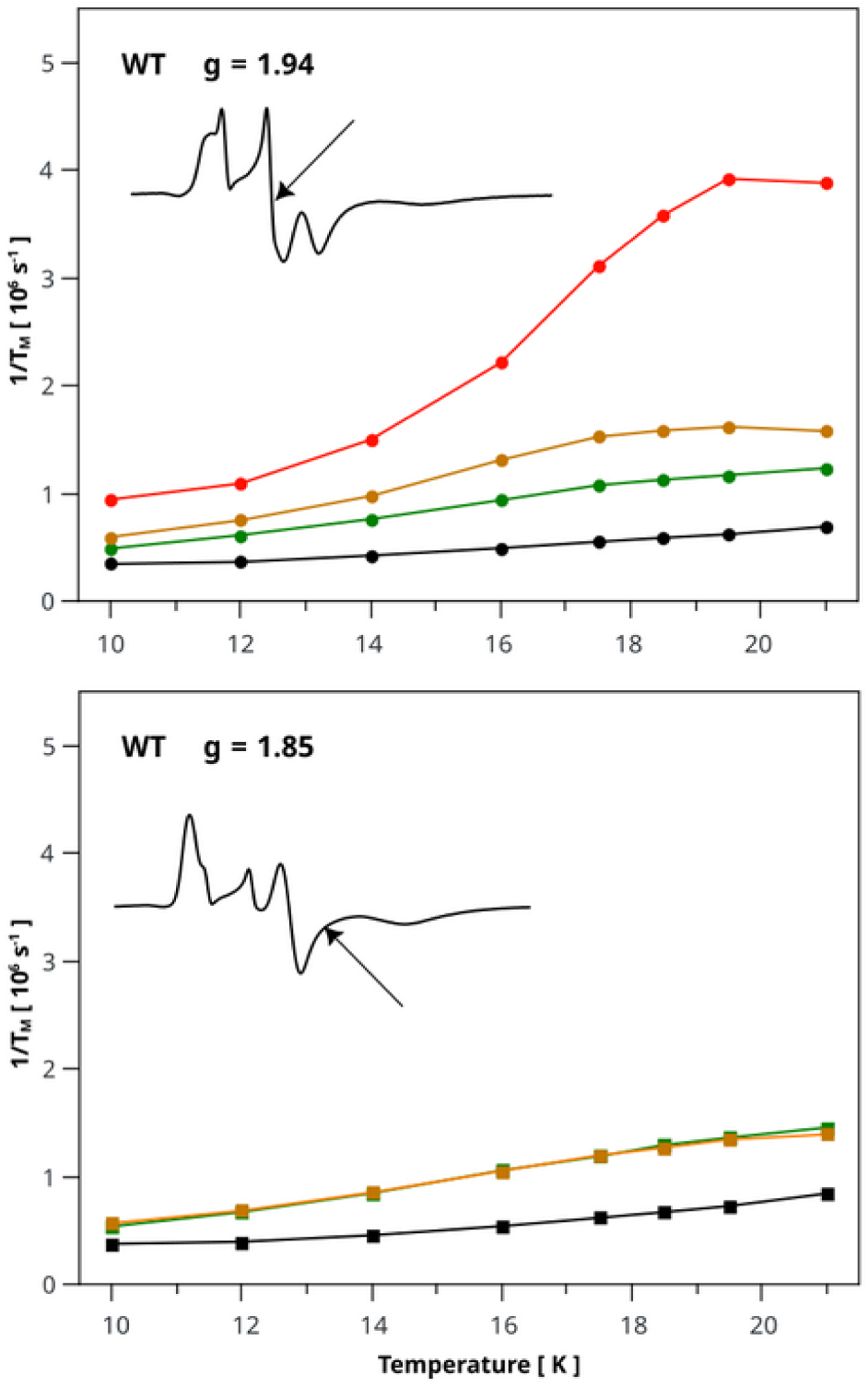
Temperature dependence of the phase relaxation rates measured for WT cyt. *bc*_1_ at the maximum of the SQ-2Fe2S^red^ signal (g = 1.94) and in the region where its contribution is minimal (g = 1.85). The rates were determined for samples frozen 3 s and 8 s after mixing with substrates (red and orange, respectively), after reaching redox equilibrium (green), and after reduction by dithionite (black).

In contrast, in the sample frozen 3 s after mixing with substrates, when heme *b*_L_ is fully oxidized and SQ–2Fe2S^red^ represents nearly 50 % of the total cluster signal, the relaxation rates increase strongly with temperature and reach a maximum of 3.91 × 10^6^ s^−1^ at 19.5 K (Fig. 4 red).

When the sample was frozen 8 s after mixing with substrates, the relaxation rates decreased substantially (Fig. 4, orange). This decrease reflects the lower fraction of SQ–2Fe2S^red^ (11 vs. 47 %) in comparison to 2Fe2S^red^ rather than a change in the redox state of heme *b*_L_ (it remains oxidized, see g_z_ transitions of heme *b*_L_ in Fig. 2). Moreover, motion of the ISP-HD may also influence the measured relaxation rate, since motion of ISP-HD away from the Q_o_ site increases the cluster–heme distance. After the sample reached redox equilibrium, SQ–2Fe2S^red^ was no longer detectable, and the relaxation rates of 2Fe2S^red^ decreased further to the levels determined by distance distribution of ISP-HD in WT. (Fig. 4, green)

At the *g* position where SQ–2Fe2S^red^ does not contribute, the relaxation rates for the sample frozen after 8 s and for the sample after reaching the equilibrium were indistinguishable (Fig. 4, bottom, green and orange) and reflects dipolar interactions between heme *b*_L_ and 2Fe2S^red^. This means that the distribution of ISP-HD in the fraction of cyt. *bc*_1_ molecules where the EB reaction did not start or was reversed is the same as in equilibrium.

To eliminate the influence of ISP-HD positional heterogeneity on dipolar relaxation rates, pulsed EPR data from the +2Ala mutant were analyzed. Figure 5 shows the results obtained for this mutant at the same *g* values as in the WT experiments: *g* = 1.94, corresponding to the maximum of the SQ–2Fe2S^red^ signal (defined on absorption spectrum), and *g* = 1.85.

**Figure 5.**
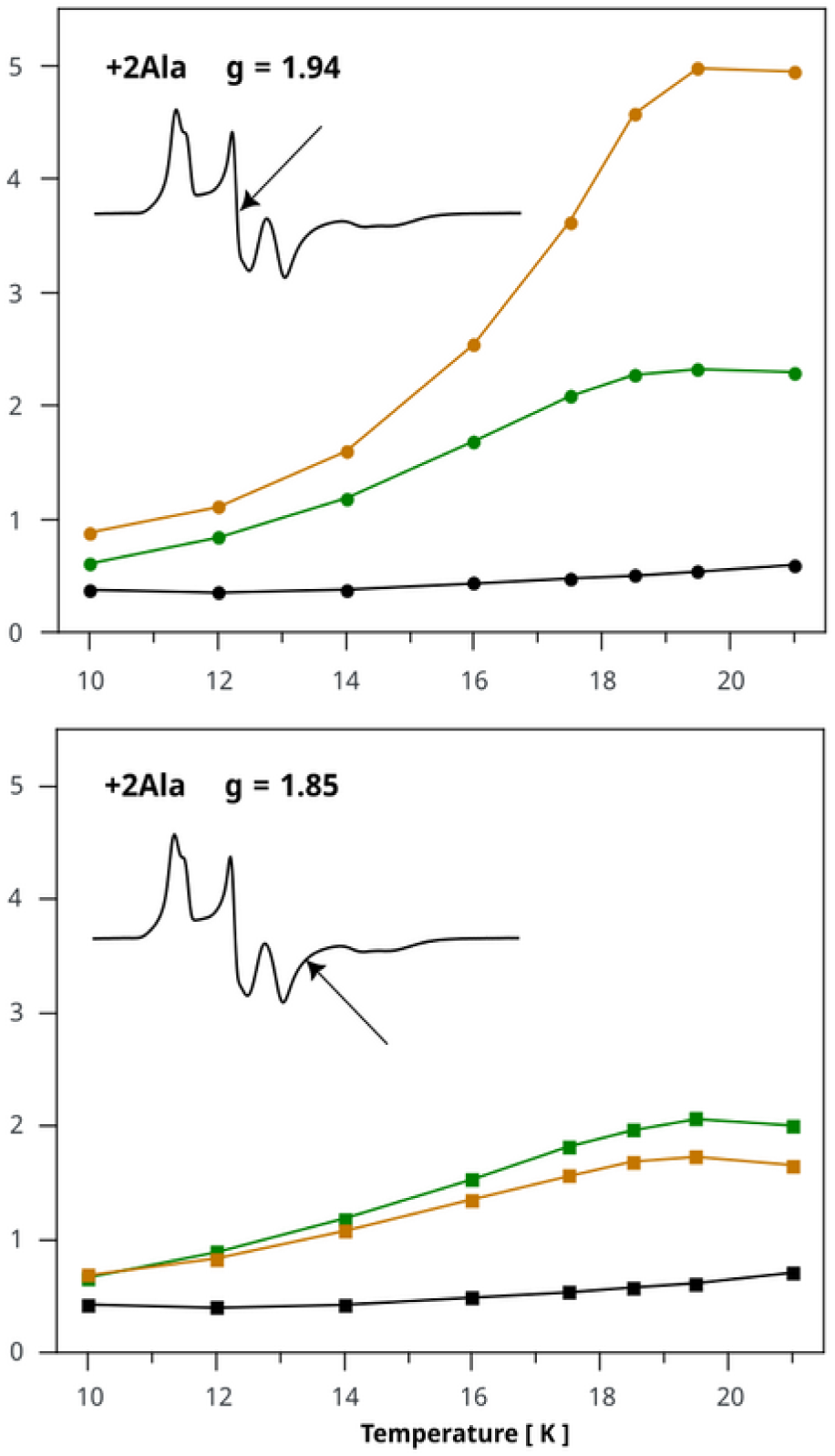
Temperature dependence of the phase relaxation rates measured for +2Ala mutant of cyt. *bc*_1_ at the maximum of the SQ-2Fe2S^red^ signal (g = 1.94) and in the region, where its contribution is minimal (g = 1.85). The rates were determined for samples 8 s after mixing with substrates (orange), after reaching redox equilibrium (green) and after reduction by dithionite (black).

For the dithionite-reduced sample, the relaxation rates were the same as in WT, indicating that arresting ISP-HD at the Q_o_ site does not affect the intrinsic phase relaxation of the cluster (Fig. 5, black). In the sample frozen 8 s after mixing with substrates, the fractions of SQ–2Fe2S^red^ and 2Fe2S^red^ (43 % and 57 %, respectively) were similar to those in WT frozen after 3 s. This allowed a direct comparison of the effect of ISP-HD motion on the dipolar relaxation rates.

In the +2Ala mutant we observed a strong temperature dependence, with a maximum relaxation rate of 4.97 × 10^6^ s^−1^ at ∼19.5 K. This value is significantly higher than in WT frozen after 3 s, despite identical oxidation levels of oxidized heme *b*_L_ and nearly identical SQ–2Fe2S^red^/2Fe2S^red^ ratios. The slower relaxation rate in WT therefore arises from the fraction of 2Fe2S^red^ that is not interacting with SQ and resides in a position remote from the Q_o_ site. In contrast, in +2Ala the ISP-HD domain remains close to the site, keeping the cluster in the shortest possible distance to heme *b*_L_. This interpretation is consistent with CW EPR spectra: in WT, the 2Fe2S^red^ signal shows no interaction with quinone at the Q_o_ site (g_x_ = 1.76) since ISP-HD occupies predominantly the remote position, while in +2Ala such interaction is detectable (g_x_ = 1.80 for Q and 1.77 for QH_2_ at the Q_o_ site, respectively).

This effect of the ISP-HD position is even more evident under equilibrium conditions. Here, SQ–2Fe2S^red^ is not detectable, and the entire phase relaxation enhancement arises from interaction between heme *b*_L_ and 2Fe2S^red^ fully bound at the Q_o_ site (Fig. 5, green). Under these conditions, the difference between WT and +2Ala reflects solely the changes in distribution of the positions of ISP-HD: in WT the cluster is largely displaced from the Q_o_ site, whereas in +2Ala the entire population of 2Fe2S^red^ remains close to heme *b*_L_. As a result, the average distance between 2Fe2S^red^ and the heme *b*_L_ iron ion is shorter.

Similar differences in relaxation rates were observed at *g* = 1.85, where the rates in +2Ala were higher than in WT (Fig. 5 bottom and 4 bottom, respectively). The small difference between the relaxation rates for +2Ala at *g* = 1.85 in samples at equilibrium and those frozen 8 s after mixing with substrates can be attributed to an additional contribution from partially oxidized heme *b*_H_ (∼25 %). Because the average distance between 2Fe2S^red^ and heme *b*_H_ is shorter in +2Ala than in WT, its influence is also reflected in the measured relaxation rates of 2Fe2S^red^ in +2Ala. In WT the effect of heme *b*_H_ is expected to be much smaller or negligible.

### Estimation of distance between SQ-2Fe2S^red^ and heme b_L_ iron ion

The lack of ISP-HD positional heterogeneity in the +2Ala mutant enabled estimation of the effective SQ–2Fe2S^red^ heme *b*_L_ iron distance based on the measured relaxation rates. The dipolar relaxation enhancement produced by a fast-relaxing center (here, the heme iron) on the phase relaxation of a slow-relaxing observed center (SQ–2Fe2S^red^ or 2Fe2S^red^) depends on the dipolar splitting, the relaxation rate of the fast-relaxing species, and the third power of the distance (*r^−^*^3^) between them. Using relaxation rates measured at the same temperature (19.5 K) and assuming similar dipolar interactions for SQ–2Fe2S^red^ and 2Fe2S^red^, we estimated the effective distance using a simplified relation between distances and relaxation rates, previously proposed for spin–lattice relaxation [72]:

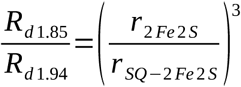

where *R*_d1.85_ is the dipolar relaxation rate measured at *g* = 1.85 (for 2Fe2S^red^), *R*_d1.94_ is the rate at *g* = 1.94 (for SQ–2Fe2S^red^), *r*_2Fe2S_ is the crystallographic distance between the heme *b*_L_ iron and the center of the 2Fe2S^red^ cluster (2.76 nm, PDB 1ZRT), and *r*_SQ−2Fe2S_ is the effective distance to SQ–2Fe2S^red^.

Based on CW EPR data, the SQ–2Fe2S^red^ signal at *g =* 1.94 is 3.6 times larger than that of 2Fe2S^red^. Thus, we fitted the ESE decay curve using a sum of two weighted exponentials: one with the relaxation rate for 2Fe2S^red^ (measured for +2Ala at equilibrium) and the second corresponding to SQ–2Fe2S^red^. This approach removed the 2Fe2S^red^ contribution from the estimated *R*_1.94_. In this case, we obtained *R*_1.94_ = 9.68 × 10^6^ s⁻¹ and *R*_d1.94_ was calculated as:

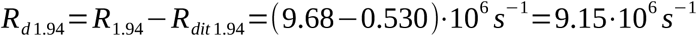

Using the reference value of dipolar relaxation rate *R*_d1.85_ for 2Fe2S^red^ calculated as:

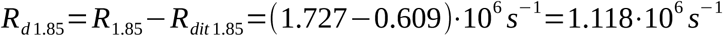

we obtained *r*_SQ-2Fe2Sred_ = ∼ 1.4 nm.

These rough estimates agree well with the distances between the heme *b*_L_ iron ion and the atoms of the stigmatellin chromone ring in the *R. capsulatus* cyt *bc*_1_ structure (1.45–1.95 nm, PDB 1ZRT) [58]. Such distances are expected if the electron spin is exchanged between 2Fe2S^red^ and SQ bound in a position similar to that of stigmatellin. Although the unpaired spin density is likely distributed over both the SQ and 2Fe2S^red^ centers, the estimated distances suggest that the dipolar relaxation enhancement of the SQ–2Fe2S^red^ pair arises predominantly from the heme iron and the nearest atoms of the spin-coupled centers.

### Interpretation of quantities of Q_o_ sites occupied by SQ-2Fe2S^red^

When samples were frozen shortly after mixing with substrates (3 s for WT and 8 s for +2Ala), we detected SQ–2Fe2S^red^ and uncoupled 2Fe2S^red^ in approximately a 1:1 ratio. This matches the full reduction of heme *b*_H_ expected under antimycin inhibition of the Q_i_ site. Importantly, the data indicate that the SQ–2Fe2S^red^ intermediate occupies, on average, only about half of the Q_o_ sites in the population; larger fractions seem not reachable. At the same time, heme *b*_L_ remained only partially reduced, reaching up to ∼11 % in WT.

Although the coexistence of SQ–2Fe2S^red^ with oxidized heme *b*_L_ had been reported previously, the fraction of SQ–2Fe2S^red^ detected earlier, using less concentrated samples, was small (typically ∼10 %). This level is comparable to the uncertainty (10 – 20 %) in determining the reduction level of heme *b*_L_ by EPR [48–52]. Thus, one could not exclude a possibility that the fraction of SQ–2Fe2S^red^ originated from the subset of monomers in which heme *b*_L_ was in fact in the reduced state. In the present work, the substantially higher concentrations of cyt. *bc*_1_ and substrates allow us to accurately quantify the involved species (*b*_L_, 2Fe2S and SQ-2Fe2S^red^). We observe that the fraction of monomers containing SQ–2Fe2S^red^ is several-fold larger than the fraction with reduced heme *b*_L_. This now clearly establishes that SQ-2Fe2S^red^ forms in monomers where heme *b*_L_ remains oxidized.

This conclusion drawn from CW EPR spectra analysis is also supported by the pulse EPR measurements, which show that SQ-2Fe2S^red^ species has much stronger dipolar relaxation enhancement than 2Fe2S^red^ in both WT and +2Ala mutant. After elimination of the influence from the ISP-HD motion we estimated the distance between the SQ-2Fe2S^red^ and iron ion of heme *b*_L_. The results clearly indicate that SQ-2Fe2S^red^ is located much closer to the heme *b*_L_ than 2Fe2S^red^, as expected. The spin echo dephasing rate is therefore dominated by the spin-coupled paramagnetic center located closest to the heme, i.e., SQ. Interestingly, the estimated distance, using simple point-dipol approximation shows that SQ is located at the distance that correspond to the position of atoms of stigmatellin bound at the Q_o_ site (PDB code: 1ZRT). This is strong experimental evidence that SQ spin-coupled to the reduced cluster is present at the Q_o_ site of the same monomers for which heme *b*_L_ is oxidized.

These observations raise two key questions. First, why does the semiquinone not reduce heme *b*_L_? Second, if all *b*_H_ hemes are reduced while all *b*_L_ hemes remain oxidized, why does only half of the dimer population form SQ–2Fe2S^red^? Addressing these questions provides important insight into the electronic configurations and cofactor interactions during active catalysis.

### Electron distribution between the hemes, quinone and the cluster

In this section, we address the first question. The observed experimental fact is that SQ-2Fe2S^red^ can only be stabilized if heme *b*_H_ is reduced. In other words, the energy of the state in which heme *b*_H_ is reduced, *b*_L_ oxidized and SQ is spin-coupled to 2Fe2S^red^ is possibly the lowest. We performed QM calculations to consider the relative energetic levels of the 3 possible configurations of redox state of the cofactors in the closest proximity to the Q_o_ site. Fig. 6 presents the energy levels for the following configurations: A) both hemes *b*_H_ and *b*_L_ reduced, Q at the Q_o_ site and 2Fe2S^red^; B) heme *b*_H_ reduced, *b*_L_ oxidized (spin β), SQ (spin α) and 2Fe2S^red^ (spin α); C) hemes *b*_L_ and *b*_H_ both reduced, SQ (spin α) and 2Fe2S^red^ (spin α). Comparison of the energy estimated for these three configurations shows that the lowest energy (0.0 by definition) is obtained for the configuration B. Configuration A has a higher energy than configuration B while C is even higher than configuration A (fig. 6 D). These results seem consistent with the experimental signatures detected by EPR. If configuration A was possible, one would detect 2Fe2S^red^ spectrum along with a flat line in the region of g_z_ transitions of the hemes *b*, which is clearly not the case in our experiments. If configuration C was possible one would detect SQ-2Fe2S^red^ signal but neither g_z_ transitions of oxidized hemes *b* nor relaxation enhancement of SQ-2Fe2S^red^ induced by the iron ion of heme *b*_L_ would be observed. In the configuration B, the expected EPR signature is the presence of SQ-2Fe2S^red^ along with the g_z_ transition for heme *b*_L_ and no transition for heme *b*_H_. This state is clearly detectable in both WT and the +2Ala mutant of cyt. *bc*_1_ under turnover conditions.

**Figure 6.**
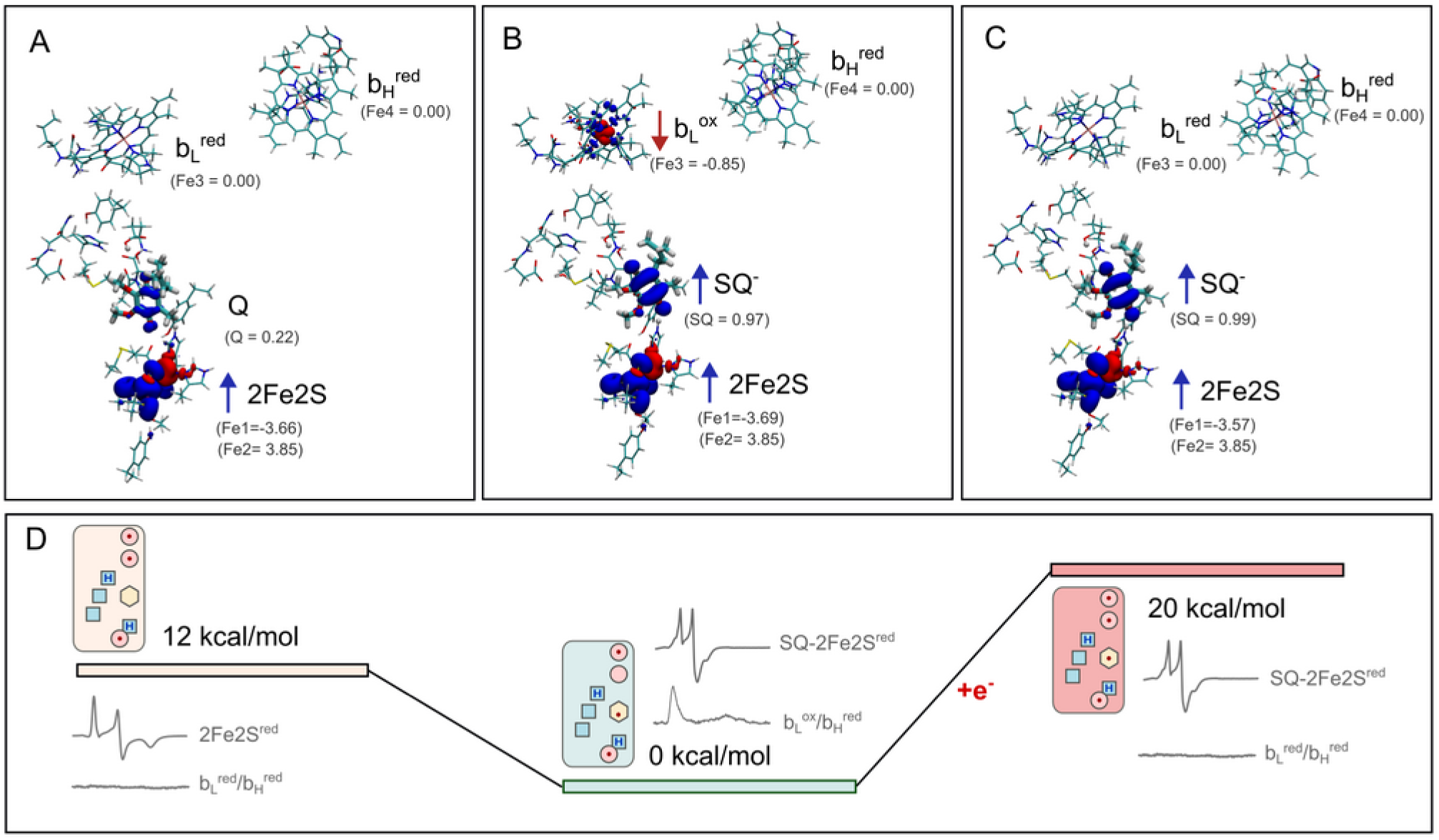
Estimation of the energy levels for three different electron configurations of cyt *bc*₁ under conditions corresponding to the reduced *b*_H_ heme, as determined by QM calculations. **A)** Configuration containing both *b*_L_ and *b*_H_ hemes, with the 2Fe2S^red^ and quinone bound at the Q_o_ site. **B)** Configuration with the same total number of electrons as in (A), but with *b*_L_ oxidized and the semiquinone (SQ) spin-coupled to 2Fe2S^red^. **C)** Configuration obtained by adding one additional electron to (B), resulting in a reduced *b*_L_ heme. Blue and red isosurfaces represent α- and β-spin densities, respectively, while the values in parentheses indicate the corresponding spin populations. **D)** Energy-level scheme illustrating the relative energies of configurations (A)–(C). The small icons represent simplified models of the respective states, where red ovals denote the *b*_H_, *b*_L_, and 2Fe2S cofactors (from top to bottom), blue squares indicate amino acid residues (E295, H276, D278) in protonated (labeled “H”) or deprotonated (empty) states (from to to bottom, respectively). Yellow hexagon denotes quinone (Q) or semiquinone (SQ). Red dots indicate the presence of an unpaired electron on a given cofactor or on SQ. Gray traces schematically represent expected experimental EPR signatures corresponding to the respective states, if they were populated.

We note, that the present observations pertain to the bacterial enzyme studied here. Because proton pathways and the protonation environment around the Q_o_ site vary across organisms, the energetic relationships between these states may not be identical in all members of the Rieske/b family. Comparative computational and experimental studies on cyt. *bc*_1_ from different organisms will be required to assess the extent of this variability.

### Electronic reconfiguration at the Q_o_ site in antimycin-inhibited cyt bc_1_

In this part we address the second question. According to canonical QBEB, the EB process is initiated by an uphill, one-electron oxidation of QH_2_ by the oxidized 2Fe2S cluster [28,73]. This first step is considered as rate limiting for EB and leads to the very transient and unstable semiquinone that is able to rapidly reduce the low-potential heme *b*_L_. In this scenario, the semiquinone formation in antimycin-inhibited cyt. *bc*_1_ should occur only to a minimal extent, through one-electron oxidation of QH₂ and under conditions where heme *b*_L_ is already reduced and unable to accept the electron originating from this SQ. For many years this mechanism seemed to be an unwavering paradigm and many studies showing only very weak signals of the single EPR line of semiquinone radical at g = 2 were interpreted as a prove of the transient and unstable nature of SQ [30,44–46,73]. However, to our knowledge, no direct experimental evidence has been provided demonstrating that the g ≈ 2 signal actually coexists with reduced heme *b*_L_. In fact, some reported data indicate that the g ≈ 2 species is magnetically coupled to paramagnetic (thus oxidized) heme *b*_L_, which is inconsistent with the assumptions of canonical QBEB [46].

EMET explains the origin of the SQ–2Fe2S^red^ signal. According to this mechanism EB is initiated by ET from QH_2_ to heme *b*_L_ in the presence of oxidized 2Fe2S [53]. If oxidation of heme *b*_L_ by heme *b*_H_ is prevented, either by antimycin inhibition or by reverse electron transfer from the Q_i_ site to hemes *b*, the electron originating from heme *b*_L_ can either end up on the 2Fe2S cluster, with the semiquinone stabilized by ferromagnetic coupling, or the reaction can be reversed to regenerate QH_2_.

Which of these outcomes occurs depends on the spin state that is transferred from the reduced heme *b*_L_ to the Q_o_ site. If α spin density is present on 2Fe2S^red^ and an α electron is deposited on SQ (by ET from heme *b*_L_), the ferromagnetically coupled SQ–2Fe2S^red^ state is formed. In contrast, if a β electron is transferred to Q, the resulting antiferromagnetic SQ–2Fe2S^red^ state, based on the recently proposed EMET mechanism, is unstable and leads to reduction of SQ back to QH_2_ by ET from 2Fe2S^red^ [53].

Because α and β electron transfer to Q seems equally probable, the resulting ratio of QH_2_ to SQ at the Q_o_ site should be 1:1. In other words, when heme *b*_H_ is reduced, the subsequent fate of the reaction at the Q_o_ site is determined by the stochastic spin state selected during electron transfer from heme *b*_L_ to Q. In neither case is a net amount of Q produced at the Q_o_ site.

According to the EMET mechanism, the reduction of SQ to QH_2_ should be associated with the oxidation of 2Fe2S cluster and disappearance of its EPR signal. However, in our experiments we see that 2Fe2S is in the reduced state. The most probable explanation is that in all states excluding SQ-2Fe2S^red^, the cluster equilibrate with the solution. Since buffer conditions are rather reducing, the cluster not interacting with SQ is mostly in the reduced form. Such explanation has been supported by the previous observation that rapid increase in the redox potential by injection of oxidized cyt. *c* to the solution of cyt. *bc*_1_ with pre-set SQ-2Fe2S^red^, affects only the free 2Fe2S^red^ cluster and not the cluster which is coupled to SQ [53]. This means that 2Fe2S can reach equilibrium with the buffer quite fast.

Alternatively, one could argue that the 1:1 ratio of SQ-2Fe2S^red^ to 2Fe2S^red^ could be explained assuming that only one monomer in the dimer, at the time, can initiate EB, while the second monomer remains inactive. In fact, models assuming only half-of-the-sites activity of cyt. *bc*_1_ have been proposed in literature. They usually considered the motion of ISP-HD to be the mechanistic element responsible for the alternation of EB activation between the two Q_o_ sites of the dimer [74–78]. However, since WT and +2Ala mutant produce the same ratio of SQ-2Fe2S^red^ to 2Fe2S^red^, the motion of ISP-HD is unlikely to be responsible for the observed proportion between these two species thus the reason must be different. In view of theoretical predictions of EMET mechnism of EB and the experimental results described here, we propose that stabilization of SQ-2Fe2S^red^ or reversing the reaction at the Q_o_ site back to the substrate (QH_2_) depends on the stochastic selection of spin α or β that are ultimately transferred from reduced heme *b*_L_ to 2Fe2S. The stochastic spin selection is a fundamental principle and does not depend on ISP-HD conformation. Thus the observed 1:1 proportion between SQ-2Fe2S^red^ and 2Fe2S^red^ does not change in structurally symmetric +2Ala when all hemes *b*_H_ are reduced and all hemes *b*_L_ are oxidized in a dimer.

## Conclusions

Our combined CW and pulse EPR analysis of highly concentrated cyt. *bc*_1_ *R. capsulatus* defined redox conditions under which the Qo site is occupied by the semiquinone spin-coupled to the reduced 2Fe2S cluster (SQ–2Fe2S^red^). Quantitative analysis of CW EPR spectra show the SQ–2Fe2S^red^ present along with oxidized heme *b*_L_ and reduced heme *b*_H_. The key argument for the co-existence of SQ with oxidized heme *b*_L_ lies in the observation that the SQ–2Fe2S^red^ population is several-fold larger than the fraction of reduced heme *b*_L_. Pulse EPR distance measurements place the SQ from the SQ-2Fe2S^red^ pair at the Q_o_-site position corresponding to stigmatellin, confirming its formation directly at the catalytic site.

QM calculations support these results. Among the accessible electronic configurations, that can be confronted with the experiment around the Q_o_ site, the state with reduced heme *b*_H_, oxidized heme *b*_L_, and SQ–2Fe2S^red^ has the lowest energy. This explains both the detectability and the stability of the intermediate under steady-state turnover conditions.

The observed 1:1 ratio of SQ–2Fe2S^red^ to uncoupled 2Fe2S^red^ is inconsistent with the canonical QBEB which predicts minute fraction of SQ only under conditions when heme *b*_L_ is reduced. Instead, our data support the general assumptions and predictions of EMET mechanism, in which bifurcation is initiated in the presence of oxidized 2Fe2S by ET from QH₂ to heme*b*_L_. Subsequently, if ET from heme *b*_L_ to *b*_H_ is not possible (i.e. heme *b*_H_ is already reduced), electron from heme *b*_L_ is reversed back to the Q_o_ site. Depending on whether an α- or β-spin electron originating from heme *b*_L_ is reversed, the SQ–2Fe2S^red^ pair is ferromagnetically stabilized or SQ is rapidly reduced to QH₂ [53].

Together, these findings reveal that semiquinone at the Q_o_ site of bacterial cyt. *bc*_1_ is not a marginal or unstable species but a robust, energetically favored intermediate arising from spin-dependent electronic reconfiguration within the dimeric Q_o_ site under conditions of reduced heme *b*_H_.

## Acknowledgment

This research project was funded in whole by National Science Centre, Poland, grant no. 2023/49/B/NZ1/02110 (MS). We gratefully acknowledge Polish high-performance computing infrastructure PLGrid (HPC Centers: ACK Cyfronet AGH) for providing computer facilities and support within computational grant no. PLG/2024/017732. We also acknowledge infrastructural support by Strategic Programme Excellence Initiative at Jagiellonian University - BioS PRA. For the purpose of Open Access, the author has applied a CC-BY public copyright licence to any Author Accepted Manuscript (AAM) version arising from this submission. We acknowledge the use of AI tools solely for linguistic improvements and text refinement, not for content creation or data analysis.

## Data availability statement

The experimental data supporting the findings of this study have been deposited in the RODBUK Cracow Open Research Data Repository under DOI: https://doi.org/10.57903/UJ/XWEWDS. The data are currently under embargo and will be made publicly available upon publication of this article.

## Notes

### Competing Interest Statement

The authors have declared no competing interest.

